# Extrinsic motivators drive children’s cooperation to conserve forests

**DOI:** 10.1101/2021.10.26.466023

**Authors:** Aleah Bowie, Jingzhi Tan, Wen Zhou, Philip White, Tara Stoinski, Yanjie Su, Brian Hare

**Author notes:** **Corresponding Author** Aleah Bowie, 104 Biological Sciences Building, Campus Box 90383, Durham, NC 27708-9976.

## Abstract

Forests are essential common-pool resources. It is increasingly critical to nurture a lifelong concern for forest health both locally and globally. Here, in two experiments, we demonstrate that school age children (6-18 yrs. old; N>1000;) of three nationalities (China, D. R. Congo and U.S.) do not have levels of intrinsic motivation to allow for successful cooperation in common-pool goods games requiring them to maintain a forest. We instead find that the size, timing, and certainty of receiving individual payoffs from cooperation significantly boost the odds of successful conservation efforts. We also provide evidence that the experience of playing this game increases longer term motivation to conserve forests. Results have implications for designing policy and curriculum to encourage collective action for forest conservation.

**One Sentence Summary:** Extrinsic motivation boosts concern for forests among children and adolescents in the United States, China, and the Democratic Republic of the Congo.

## Main Text

Forests are vital to human and planetary health. They are common-pool goods that require both local and international cooperation to maintain (Van Vugt, 2009). Educating the public to support forest health through personal behavior and policy remains a top priority for governments and conservationists. School aged children are often regarded as one of the most important audiences for this education. Early positive exposure to wildlife and forests is commonly believed to translate into prosocial behavior toward forests throughout life (Bowie et al, 2020). However, it remains unclear what type of educational experiences might encourage prosocial behavior toward forests across diverse populations (Saylen & Blumstein, 2011).

The *Biophilia Hypothesis (BH)* suggests humans evolved intrinsic motivation to care for the natural world (Wilson, 1984; Kellert & Wilson, 1993). It predicts that universally across cultures our selfish need to interact with life motivates humans to protect natural areas (Kahn, 1997). Early exposure to nature should nurture this intrinsic motivation and result in increased expression while extrinsic rewards may dampen it (Warneken & Tomasello, 2008; Ariely, 2009).

In contrast, the *Anthrophilia Hypothesis (AH)* posits that humans evolved intrinsic motivation for prosocial behavior towards kin, ingroup members and strangers, but not more abstract social categories like future-others or forests (Singer, 1981; Warneken et a., 2007; Silk and House, 2011; Silk, 2002; 2006; Chapais, 2001; Hill & Hurtado, 2017; Moore, 2009). Any prosociality toward abstract social categories is an accidental by-product and emergent property of plasticity in our evolved motivation to help other humans. The AH predicts that cross-cultural variability in prosociality toward forests is shaped by ecological and economic uncertainty (Eom et al., 2015; Frankenhuis et al, 2015; Mittal et al, 2014; Van der Lindgon, 2015; Spence et al., 2012; Boyd et al, 2010; Henrich et al, 2006). Experiences that incorporate extrinsic rewards and teach the link between material gain, reputation enhancement or punishment and forest conservation are likely needed for children to internalize the value of these shared resources (Ryan & Deci, 2000).

School age children provide a strong test of these hypotheses. They are old enough to understand the concept of a common-pool good (children as young as six behave strategically in similar public goods games; (Keil et al, 2017; Voglesang et al, 2014; Hermes et al, 2019; Yang et al, 2018; Koomen and Hermann, 2018; Englemann et al 2018)), but their motivations are not yet fully shaped by adult participation in economic markets. However, little, if any, experimental work has examined the willingness of children to help abstract, nonhuman entities like forests (Koomen & Hermann, 2018).

Adult motivation for conservation has been tested experimentally using a collective-risk common-pool goods game (Milinski et al, 2008). In this game, the entire group is threatened with losing their endowment unless individual donations exceed a threshold needed to maintain a common-pool good. In Western populations the certainty of personal loss, reputation, and the immediacy of the benefit the good delivers largely determine group success (Milinski et al 2008; Jacquet et al 2013; Hauser et al, 2014). What is needed to better understand the origin of these preferences is a version of this paradigm designed for cross-cultural use with children.

Previous research has shown that cognitive preferences relating to certain decision making are shaped by considerations of resources within environments (Ellis et. al 2009; Bateson et al., 2014, Belsky, 2008). Frankenhuis et al. (2015) identified two ecological factors that influenced decision-making: harshness, defined as the rates of mortality and morbidity caused by factors an individual cannot control, and unpredictability, defined as the change in mean variation in harshness over time. This framework can therefore provide explanations of populational differences in decisions about resource distribution. Populations in highly uncertain environments tend to be more vigilant, more risk prone, and steeper temporal discounters than those in less uncertain environments (Ellis et al., 2009; Salali et al., 2015; Mittal et al., 2014).

Based on Milinski et al (2008), we designed a game in which school aged children from three different countries (N>1,000; 6-12 years of age) decided what portion of an endowment they wished to contribute to maintaining the conservation of a local forest. Six age matched peers played together and received tokens after an orientation from an experimenter (Appendix S1). They learned the goal of the game was for their group to meet a donation threshold required to keep a forest healthy. Children could anonymously decide to keep the tokens they received until the end and exchange them for prizes (i.e. toys or candy) or they could contribute any portion to local forest conservation. They were told they would have a set number of trials to reach the goal. The experimenter added tokens to a *Connect Four*® board(s) after each trial within a round to display the cumulative number of tokens given over the course of the trials (Fig S1). The board allowed children of all ages to visually understand how close they were to reaching the threshold. To test the predictions of our hypotheses, in two experiments, we varied the type and amount of extrinsic motivation for donating. Donation patterns across conditions were analyzed at the group and individual level using linear regression models designed to account for age, sex, education level, nationality, and difference between the threshold number and accumulated tokens per trial. The BH predicts a universal level of intrinsic motivation across cultures that will only be reduced by extrinsic motivators. The AH predicts cooperation will be strongest in response to high extrinsic rewards or punishments and will vary cross culturally depending on the level of uncertainty characterizing a group’s environment.

In a first experiment we tested how risk of losing one’s own rewards influenced motivation for prosocial behavior towards a forest by testing children in the United States (N=198), the People’s Republic of China (N=216) and the Democratic Republic of the Congo (DRC; N=156; see Table S1-2). Children from these countries provide a powerful test of our hypotheses because they vary on country-wide levels of resource uncertainty. Based on country-level statistics of life expectancy, health outcomes, and GPD per capita (WHO, 2019), we classify the DRC as a comparatively more resource uncertain, and the USA being comparatively less resource uncertain, with China in between.

After being oriented, each group was given warm-up trials to practice the donation procedure before the test phase. Children were then assigned to one of three motivation conditions that only differed in the risk of forfeiting their earnings if the group failed to meet the donation threshold needed to care for the forest (Fig 1A):

**Figure 1.**
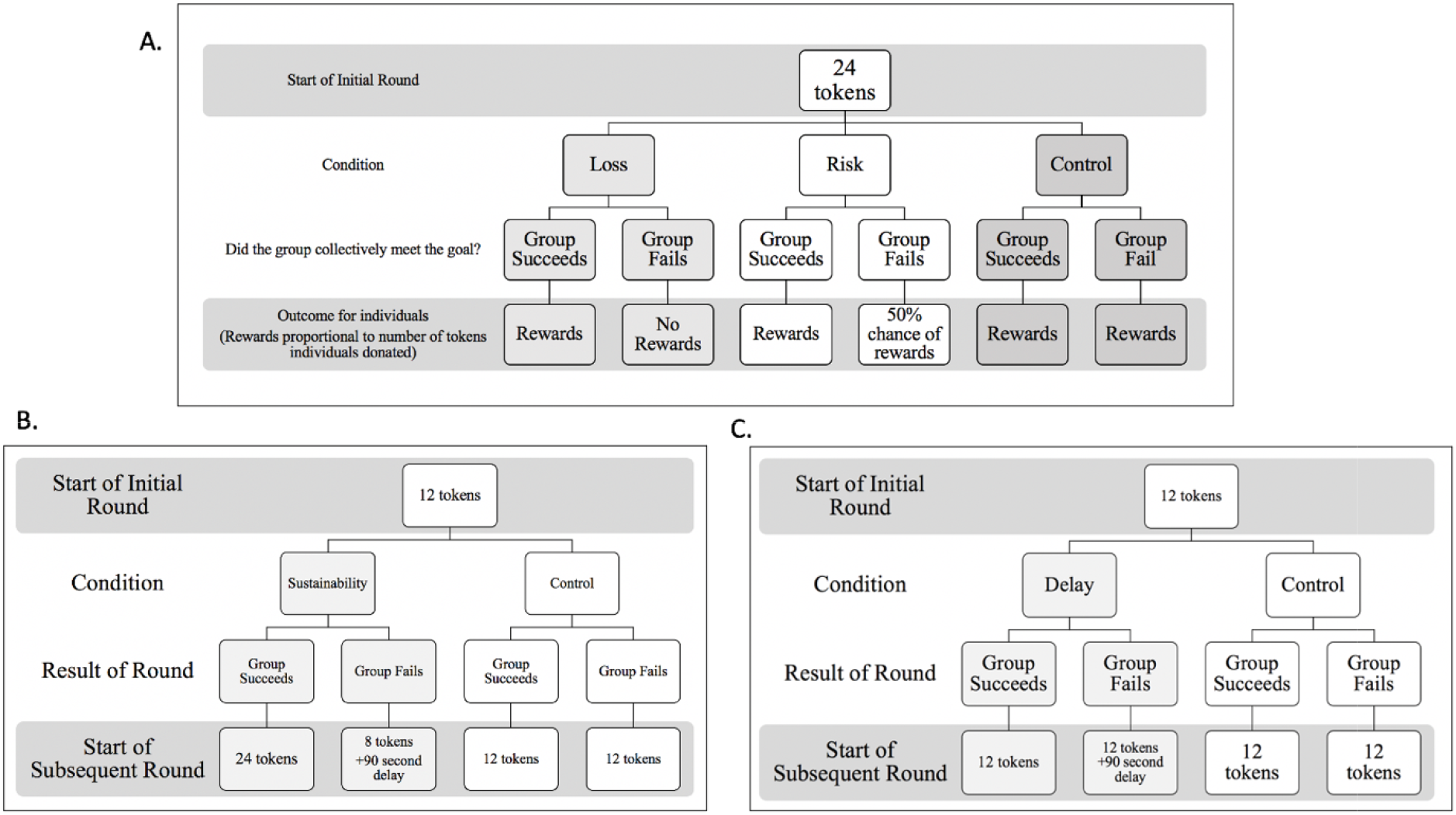
**A)** Outcomes for conditions in Experiment 1. Group Success indicates if the group collectively gives enough to the collection bank to meet the conservation goal (72 tokens). Reward indicates that individuals in the group can exchange their remaining tokens for rewards. B) Differential outcomes for the Sustainability and Control Conditions in Experiment 2 C) The Delay and Control study in Experiment 2. Success in a round for B and C required the group to collectively donate minimum of 36 tokens.

*Loss Condition* – Failure to meet the donation threshold results in all participants losing all their tokens earned in the game.

*Risk Condition* - Failure to meet the donation threshold results in a coin toss giving participants a 50% chance of keeping or losing the tokens earned in the game.

*Control Condition* - Participants keep the tokens they earn in the game regardless of whether their group meets the donation threshold.

Before starting the experiment, children received 24 tokens, learned the risk to their earnings if their group failed to meet the donation threshold, and were told they had six trials to succeed. They could donate 0, 2 or 4 tokens per trial and the donation threshold was 72 tokens per round (requiring 2 *Connect Four*® boards to display). Success required an average donation of at least 2 tokens per trial per child (6 participants X 2 tokens X 6 trials).

Using Poisson regression models, we found that extrinsic motivation led to more success meeting the threshold. While only 37.5% of groups succeeded in the control condition, 79.3% and 91.2% succeeded Risk and Loss extrinsic conditions, respectfully (Fig S3). Groups across all three nationalities gave more in the Risk [z=6.66, p<.001] and Loss condition [z=6.90, p<.001] than they gave in the Control condition (Fig 1).

Countries’ resource uncertainty was linked to donation levels. The group-level analysis demonstrates that children from DRC donated significantly less compared to Chinese children [z=-9.36, p<.001] (Fig 2) and children in the USA also gave less than Chinese children [z=-4.82,p<.001]. Children growing up in the more resource uncertain environment showed reduced donations as they aged while donations increase with age in the comparatively more resource certain countries. The individual level analysis shows that compared to Chinese children, older Congolese children gave fewer tokens than younger children [z=-1.98, p=.047] while the opposite relation was found in the U.S. sample [z=2.87, p=.004; see Fig.S5] in an age-matched analysis.

**Fig 2.**
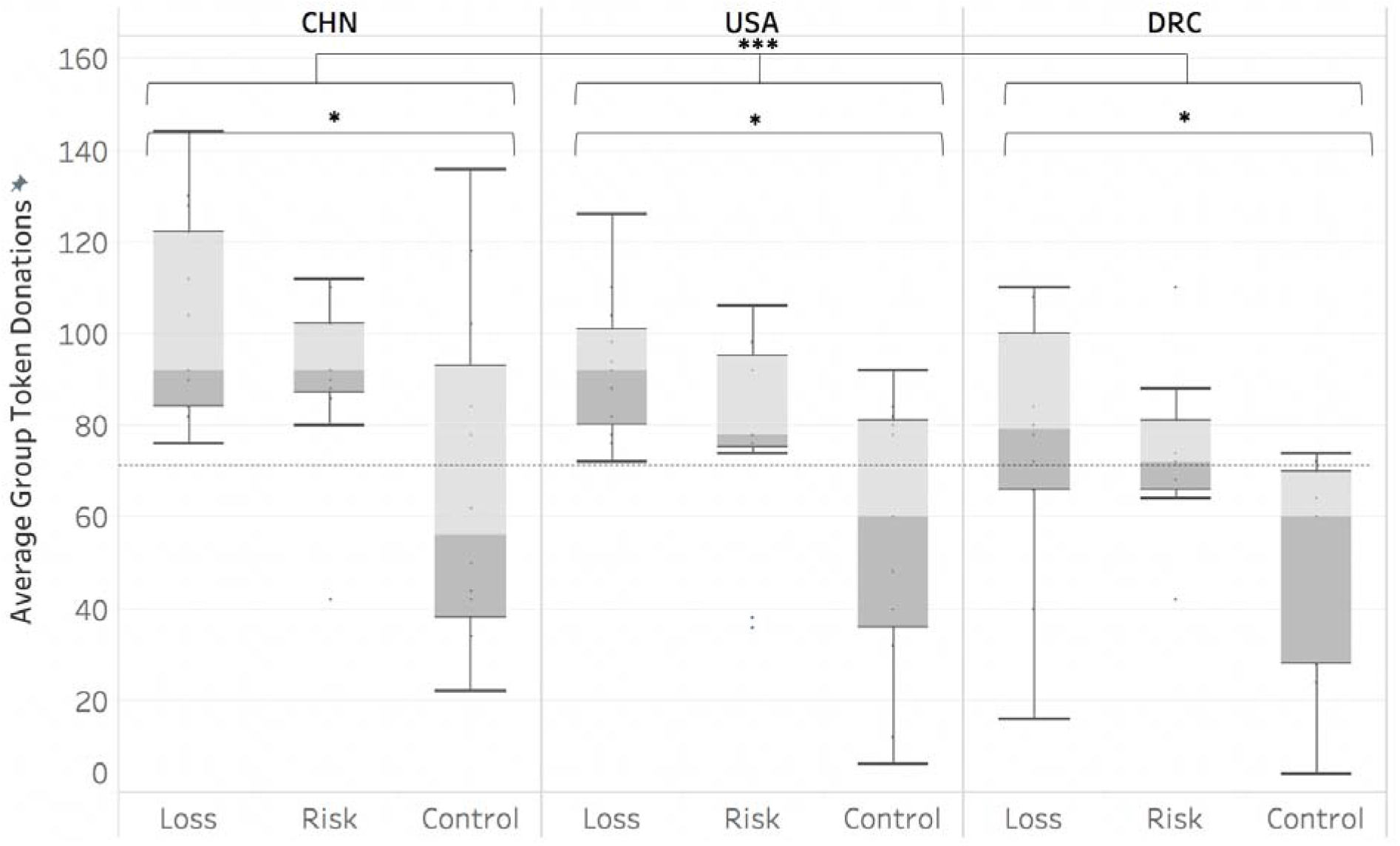
Average total group (± SD) donations by condition and country. The horizontal dashed line represents the number of tokens (72) needed to reach the conservation threshold. *p<.05

The individual level analysis also assessed individual donations levels once the threshold was met and again found little evidence for intrinsic motives, especially in the DRC and USA samples. Results detected that across conditions, individual donations dropped significantly in trials after the donation threshold was met by children from the DRC [z=-2.60, p=.009] and the US [z=-4.23, p<.001], but not in China (Fig. S6). Only children from China continued to donate to aid the forest once the donation threshold was met and their own rewards were secured (perhaps an expression of collectivist cultural norms or higher sensitivity to experimenter demand effects (Kagitcibasi, 1997)).

In a second experiment we tested children (N=516) from a summer camp at an American zoo using a variation on the aforementioned common-pool goods game (Table S3). The conditions examined the use of time delays as an extrinsic motivator as opposed to risk of losing resources as the extrinsic motivator.

*Sustainability condition* – children (N=216) participated in an instructor led discussion on the value of forest conservation right before they began the game. It was then explained that in the game their donations would be used to help keep a forest healthy. During the discussion they were told 1) the tokens they receive represent money made from selling lumber from the forest they were to manage 2) meeting the donation threshold (filling the collection bank with 36 tokens) increased the forest productivity while failing reduced it and 3) success translated into 24 tokens for everyone in the next round while failure reduced the productivity of the forest to 8 tokens per player and required a 90 second waiting period between rounds while the forest recovered on its own (Fig 1B).

*Sustainability Control Condition* – children (N=144) did not discuss forest conservation and were told they received the same 12 token endowment after each round regardless of whether the group met the threshold. There was never a delay between rounds.

*Delay Condition* – children (N=78) did not discuss forest conservation and were told failure to meet the donation threshold and fill the collection bank would result in a 90 second delay before the next round. They were told they would receive the same 12 token endowment regardless of success or failure at meeting the threshold (Fig 1C).

*Delay Control Condition* – children (N=78) did not discuss forest conservation and were told they received the same 12 token endowment after each round regardless of whether the group met the threshold. There was never a delay between rounds.

In this iteration of the study, there were three trials per round, with up to 3 rounds total. Children could donate 0,2 or 4 tokens per trial, and the donation threshold was 36 tokens per round.

Based on our linear regression, groups that experienced extrinsic motivation were far more successful meeting the donation threshold (see Table S5). Participants in the Sustainability condition contributed significantly more than groups in the Sustainability Control condition [z =5.65, p<.001] and participants in the Delay condition contributed more than those in the Delay Control condition [z=2.22, p=.026]. At the individual level, participants in the Sustainability Condition gave more tokens than those in Sustainability Control Condition [z=8.98, p<.001] and individuals in the Delay Condition gave more tokens than those in Delay Control [z=4.12, p<.001].

A significant effect of condition was detected when all four conditions were directly compared [F(3,81)=28.26, p<.001; one-way ANOVA; see Fig. 3]. Groups donated more tokens in the Sustainability and Delay conditions than in the control conditions [p<.001; Post-hoc Tukey tests]. There was no significant difference between the two control conditions or between the experimental conditions. By this measure, the threat of a delay in obtaining the endowment in future rounds was as successful as the more complete simulation of forest management in the Sustainability condition (i.e. few groups failed to meet the threshold in the Sustainability (2 out of 36 groups) as well as in the Delay conditions (4 out of 13 groups)).

**Fig 3.**
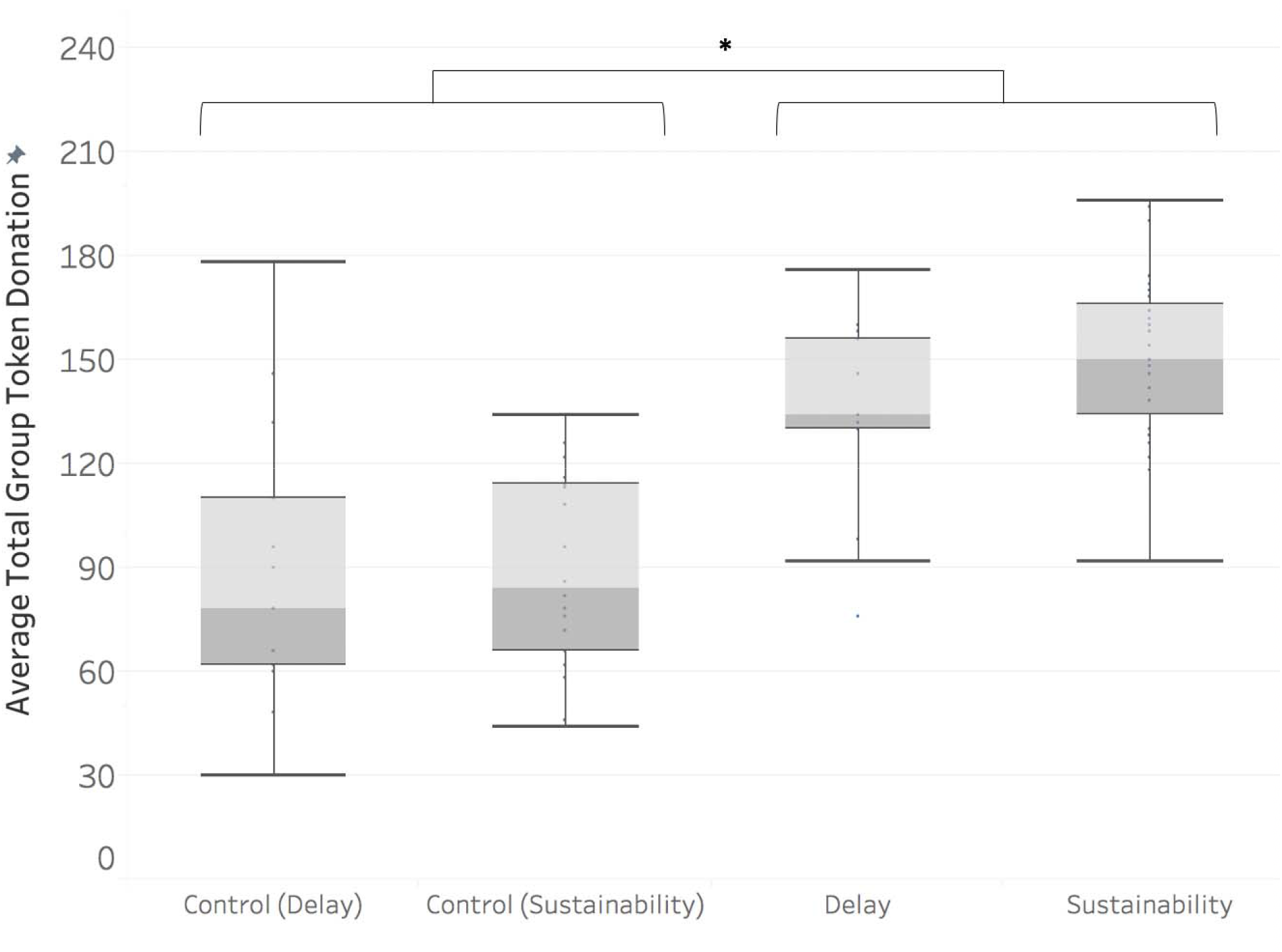
Average total group token donation (± SD) by condition in Experiment 2. *p<.05

Finally, to test the impact of participating in the Sustainability condition on motivation for forest conservation, some participants (n=36) were given an opportunity to express their opinion about the future of a local forest. These responses were made an average of 5 days (+/- .6 SE) after participating in the initial experiment and were compared to children (n=25) who attended the same camp session but had not previously participated in our experiments. The children were told the local mayor was to decide if a forest would be cleared to make money for the city and that the mayor wanted to hear their opinion. They received postcards on which they could send her their message. They were told there was no correct answer. Coders blinded to condition then quantified the effort and content of what children drew and wrote on each postcard (see Table S4).

This follow-up study found that experiencing the full model of sustainable practice resulted in internalization of motivation to conserve forests (see Table S6). Children who had participated in the Sustainability condition an average of five days earlier were more likely to make a drawing on their postcard (df=59, t=2.07, p=.04Welch t-test) and put more effort into the drawing than the control group (df=54, t=2.41, p=.019, Welch t-test). This supports the prediction that experiencing simulated forest management can increase motivation to save forests beyond the initial experience of the game.

Overall the results of our experiments support the predictions of the Anthrophilia Hypothesis. Across all three nationalities, children were most likely to successfully support forest conservation when extrinsic motivation was highest. Even children attending an environmental camp in the U.S. were unlikely to meet the donation threshold without rewards or punishment. Cultural differences and differences in population level resource certainty likely also shape donation preferences in this simulation. In the second experiment, we demonstrated that even minimal extrinsic motivation boosted cooperation--just the threat of minor time delay between rounds significantly increased success in reaching the threshold in children of all ages. The more effortful drawing behavior on postcards of children who participated in the most complete modification if this common-pool goods game suggests this type of experience has potential to drive longer term behavior change.

There was limited evidence in support of the Biophilia Hypothesis. Intrinsic motives were not strong enough to consistently drive cooperation across nationality or condition. Only a minority of groups succeeded when there was no personal consequence associated with group success or failure. When failure resulted in loss of earnings, children in the US and DRC curtailed donations as soon as their own selfish rewards were secured. This suggests children understood how to maximize their individual payoff, but they were only motivated to give minimally when the common-pool good conflicted with their own interest. The exception to this pattern were Chinese children who donated even after the threshold was met.

These findings suggest that in addition to experiencing wildlife, children benefit from experiencing the decisions required to protect common-pool goods to develop more lasting concern for forest conservation. Future research can explore how to translate common-pool goods games into curricula. Cross-cultural research is also badly needed, especially with children from developing countries with more resource uncertainty or different cultural norms. With such knowledge, conservation education will be far more effective. Vital common-pool goods – including forests – will experience enhanced protection. Both people and wild places will benefit.

## Acknowledgments

The authors wish to thank all the participants, the staff of Lola ya Bonobo, Fanny Minesi, Claudine André, Raphael Belais for continuous support, Blaise Mwaki for data collection in the DRC. The Yongding Branch of Beijing No. 2 Experimental Primary School helped with data collection in China. Thanks also to support at Zoo Atlanta from Michelle Kolar, Staci Wiech, and Yeta Robinson, Daniel Hontz, and Zoo ATL research assistances Amanda Danner, Feruthe Kidane, Gianna Ossello, Julia Villegas, and Kyle Smith.

## Data Availability Statement

The data that supports the findings of this study are available in the supplementary material of this article.

## Ethics

Ethics approval for all studies was granted by Duke University Campus IRB protocol #2017-1004 (USA and DRC) and protocol # 2017-1054 (China).

## Author Contributions

AB, JT, WZ, BH designed the study, AB and PW analyzed the data. AB and BH wrote the manuscript. TS, YS and provided resources and supported study implementation.

## Supplementary Materials

Materials and Methods

Supplemental Analysis

Figures S1-S9

Tables S1-S5

Appendix S1

External Database S1

